# Environmental DNA sequencing data from algal blooms in Lake Erie using Oxford Nanopore MinION

**DOI:** 10.1101/2022.03.12.483776

**Authors:** Alexander F. Koeppel, Will Goodrum, Morgan Steffen, Louie Wurch, Stephen D. Turner

## Abstract

Harmful Algal Blooms (HABs) pose a significant and increasing risk, both to human health and to the Blue Economy. Genomics approaches to early detection promise to help mitigate these risks. We have developed and prototyped HABSSED (HAB Surveillance by Sequencing of Environmental DNA), a portable, reliable, rapid, low-cost pipeline for detecting HABs in the field using 3rd generation sequencing with the Oxford Nanopore MinION device. We demonstrated the efficacy of our approach by sequencing existing samples collected from a National Oceanic and Atmospheric Administration (NOAA) Great Lakes Environmental Research Laboratory (GLERL) on Lake Erie. We sequenced environmental DNA (eDNA) from samples drawn before, during, and after a *Microcystis* bloom, and estimate the abundanced of HAB-associated taxa. While sequencing results showed some evidence of human and *E. coli* contamination, we find that the abundance of *Mycrocystis* and other HAB-associated orgnisms significantly differs between pre- and post-bloom environments. Here we describe the publicly available sequencing data that was generated as part of this research, which is available in the Sequence Read Archive (SRA) under accession number PRJNA812770.

## 1 Background & Summary

Harmful algal blooms (HABs) represent a significant challenge to the health of the world’s waters and the Blue Economies that depend on them. Toxic HABs of many different species represent a direct risk of injury or death to humans (Erdner et al. 2008; Jochimsen et al. 1998), but even algal blooms that are not directly toxic can cause significant damage by leading to hypoxia or anoxia in the water column, shading critical submerged aquatic vegetation, and degrading ecosystem function (Heisler et al. 2008). As a result, HABs cause significant harm to the Blue Economy, adversely affect beaches (Morgan, Larkin, and Adams 2009) and fisheries (Holland and Leonard 2020), and cost hundreds of millions of dollars in efforts to manage and mitigate the blooms (Anderson et al. 2008). As the global climate continues to warm, the number and severity of HABs is expected to increase, along with their incumbent deleterious health and economic effects (Anderson, Glibert, and Burkholder 2002; Fu, Tatters, and Hutchins 2012; Moore et al. 2008).

We are developing a genomics-based approach to help the diverse group of Blue Economy stakeholders to address this challenge. We developed a prototype of HABSSED (Harmful Algal Bloom Surveillance by Sequencing of Environmental DNA), an environmental DNA (eDNA) sequencing and bioinformatic analysis pipeline for detection of toxic *Microcystis* blooms using third generation sequencing technology, specifically Oxford Nanopore’s MinION device. MinION sequencing technology has numerous advantages over more traditional next generation sequencing platforms (e.g. Illumina HiSeq and/or MiSeq) that make it an ideal choice for HAB monitoring. The MinION sequencer is handheld and robust to vibration and other movement, can be powered by the same laptop used for data analysis, and has a relatively low initial capital investment for the instrument, allowing it to be used in field environments and putting the technology within reach of even small labs. In addition, the MinION is capable of truly de novo sequencing, as no sequence-specific primers are needed; any DNA within the sample can be bound to sequencing adapters and processed. As single molecules of DNA are sequenced, this data is accessible to the user in nearly real-time; analysis can begin as soon as five minutes after sequencing starts, rather than the 24+ hours of a conventional NGS sequencing run. For detection purposes, samples can be effectively sequentially multiplexed onto a single flow cell, decreasing operational costs.

The ultimate goal of the HABSSED is to include all steps necessary for research and monitoring personnel to rapidly obtain an estimate of aquatic microbial species abundance for HAB-related organisms from a raw water sample (Figure 1). HABSSED is being designed to be field-portable, incorporating genomics tools and workflows that can be run on a well-powered laptop or using an Amazon Web Services (AWS) or other similar cloud instance. Ultimately, field researchers or monitoring professionals will be able to readily detect the presence and abundance of toxic *Microcystis* with minimal specialized training, even aboard ships or in other remote field locations.

**Figure 1:**
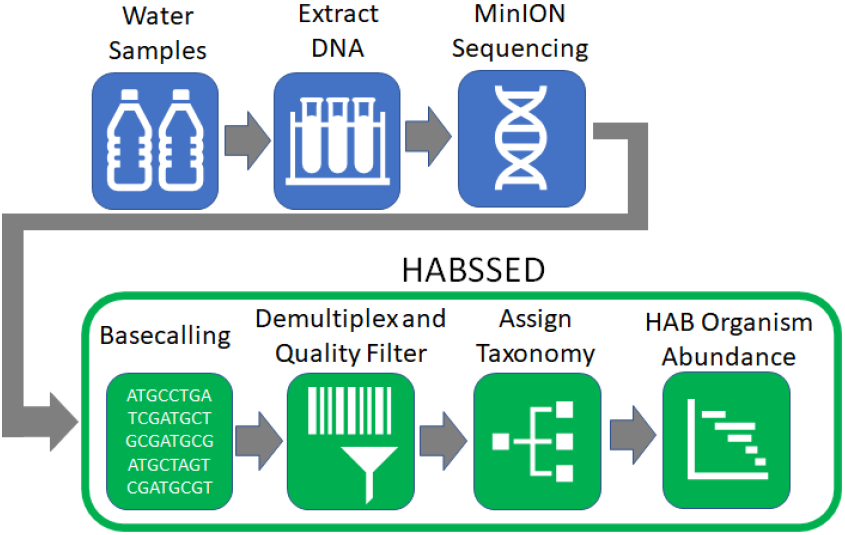
The HABSSED pipeline. eDNA is extracted from water samples and sequenced using the MinION. Bases are called and reads are demultiplexed and filtered for quality. Taxonomy of reads are classified, and abundances of each taxon is calculated based on read counts.

For developing and evaluating the HABSSED capability we used MinION to sequence ten samples from several NOAA Great Lakes Environmental Research Laboratories (GLERLs). Here we describe the methods used for sample collection, sequencing, and data deposited to the Sequence Read Archive (SRA) under accession number PRJNA812770.

## 2 Methods

### 2.1 Sample collection

We collected water samples from NOAA GLERL stations WE02 and WE13 during a 2019 *Mycrocystis* bloom in Lake Erie. Collection sites are show in the map in Figure 2 (adapted from www.glerl.noaa.gov). Sampling excursions from these NOAA GLERL stations in the 2019 bloom season took place during pre-, peak, and post-bloom conditions. Samples were drawn using standard collection protocols (Steffen et al. 2017) at a water depth of 0.5m-1.5m. Metadata on the physical and chemical properties of the water at the time of sampling were also collected (Boedecker et al. 2020).

**Figure 2:**
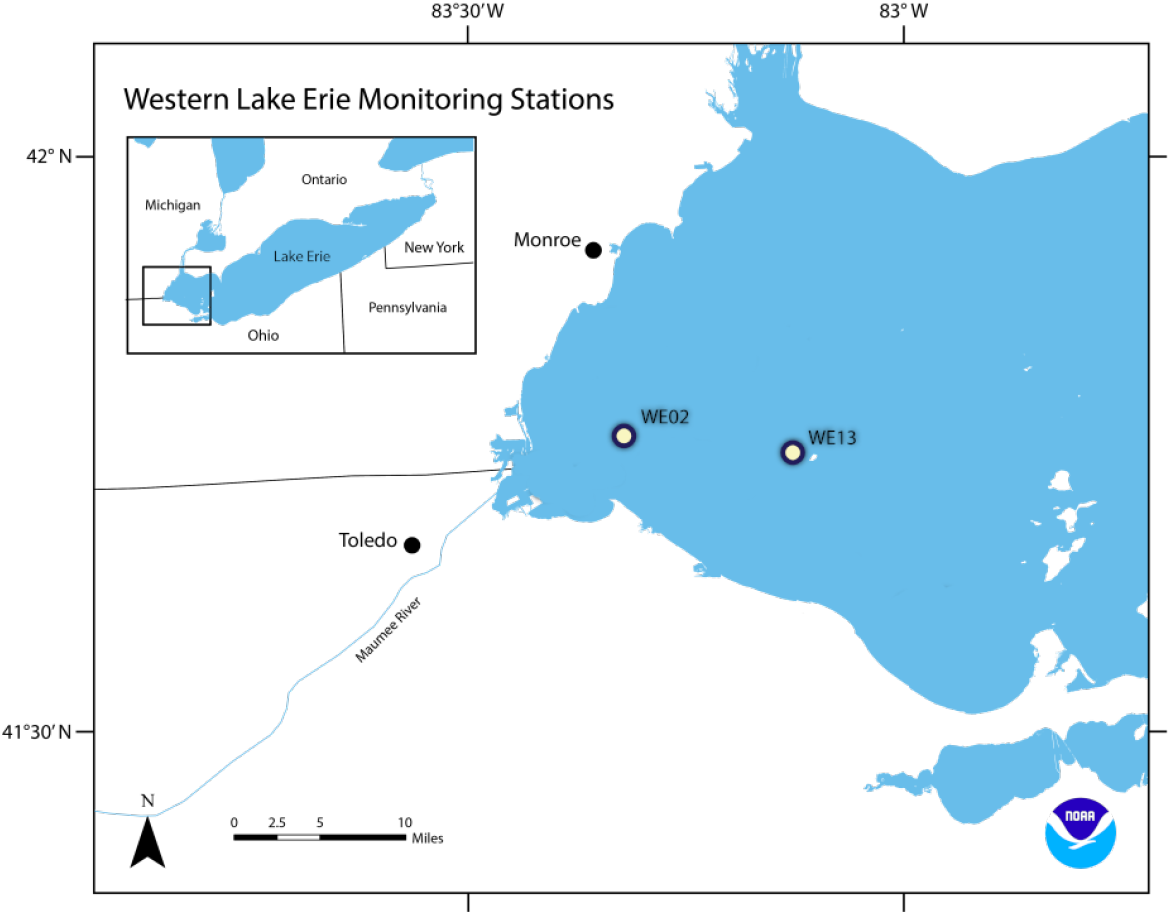
GLERL sites WE02 and WE13 from which water samples sequenced here were collected (adapted from https://www.glerl.noaa.gov/).

### 2.2 Sequencing and data analysis

Biomass was collected onto 0.2 um Sterivex filters and kept on ice until they were returned to the lab where they were stored at −80C until extraction. Extraction protocol was adapted according to (Cruaud et al. 2017). We then prepared the libraries for MinION sequencing using Oxford Nanopore Technologies (ONT) sequencing kit (initially the Rapid Sequencing Kit (SQK-RAD004) but transitioning to the Rapid Barcoding Kit 96 (SQK-RBK110.96) for later runs). We sequenced the eDNA from those samples using the MinION Mk1C device, which performed base-calling and quality filtering using ONT’s embedded MinKNOW software using the default settings. We then performed taxonomic classification using the ONT What’s In My Pot (WIMP) pipeline and counted the abundance of *Microcystis* reads, as well as species and strains within the *Microcystis* genus where possible. We computed the relative abundance by dividing these counts by the total number of reads passing the QC filters for the same samples.

## 3 Data Records

All sequencing data is available in the Sequence Read Archive (SRA) at accession PRJNA812770. Table 1 provides a subset of the metadata associated with these samples. The full metadata is available on the SRA bioproject page.

**Table 1:** Subset of the available metadata about samples sequenced in this study, available at SRA accession number PRJNA812770.

| Sample | Run date | GLERL | Condition | Input (ng/uL) | Run type | # Multiplexed |
| --- | --- | --- | --- | --- | --- | --- |
| FR1_WE13_PB6 | 2021-10-07 | WE13 | Pre-bloom | 7.98 | Single sample | 1 |
| FR2_WE02_PB2 | 2021-11-08 | WE02 | Pre-bloom | 0.353 | Single sample | 1 |
| MP1_RB1 | 2022-01-26 | – | Reagent blank | LOW | Multiplexed | 5 |
| MP1_WE02_PB1 | 2022-01-26 | WE02 | Pre-bloom | 0.264 | Multiplexed | 5 |
| MP1_WE13_PB1 | 2022-01-26 | WE13 | Pre-bloom | 0.053 | Multiplexed | 5 |
| MP1_WE02_B1 | 2022-01-26 | WE02 | Bloom | 22.9 | Multiplexed | 5 |
| MP1_WE13_B1 | 2022-01-26 | WE13 | Bloom | 12.4 | Multiplexed | 5 |
| MP2_RB1 | 2022-02-16 | – | Reagent blank | LOW | Multiplexed | 6 |
| MP2_NC1 | 2022-02-16 | – | Negative control | LOW | Multiplexed | 6 |
| MP2_WE02_PB2 | 2022-02-16 | WE02 | Pre-bloom | 0.159 | Multiplexed | 6 |
| MP2_WE13_PB2 | 2022-02-16 | WE13 | Pre-bloom | LOW | Multiplexed | 6 |
| MP2_WE02_B2 | 2022-02-16 | WE02 | Bloom | 6.16 | Multiplexed | 6 |
| MP2_WE13_B2 | 2022-02-16 | WE13 | Bloom | 10.8 | Multiplexed | 6 |

**Table 2:**
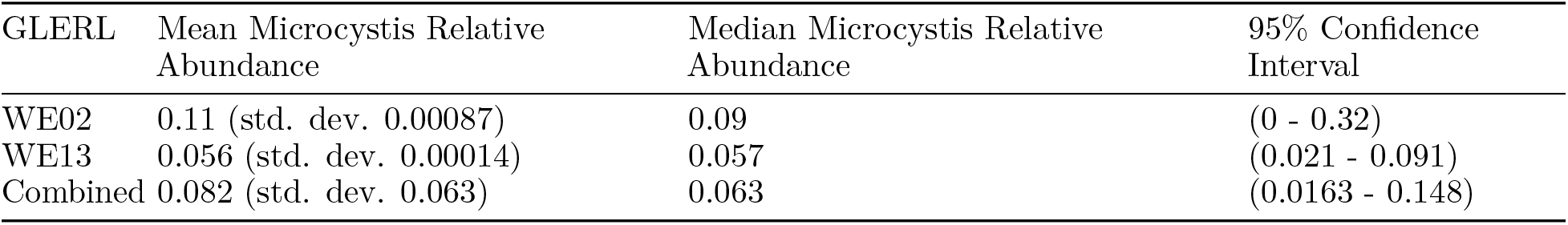
Average abundance of *Microcystis* between stations and across samples.

## 4 Technical Validation

We found that the relative abundance values are comparable between samples and sampling stations (GLERLs). Though the abundance does appear to be higher at GLERL WE02, this difference is not statistically significant due to the wide range of variation within the samples from each GLERL (t-test, *p* = 0.407). This resulted a broad baseline range of pre-bloom *Microcystis* abundance, ranging over almost a full order of magnitude from 0.016% to 0.15% of reads in a given sample.

Additionally, there was notable human and *E. coli* contamination present in several samples that was not revealed until taxonomic classification. While we took steps to track down and eliminate the contamination for later samples, we were unable to conclusively discover the source. Based on the quantities observed in the negative controls and reagent blanks, these included at minimum, 1,400 to several thousand reads classifying as *E. coli*, and from 50 to several hundred reads classifying as *H. sapiens*, and hundreds more classifying to other taxa, including *Shigella*, *Acinetobacter*, and *Microcystis* (though never more than 10 reads in any of the blanks or negative control). In some of the bloom-drawn samples (those from GLERL WE13) the abundance of *E. coli* was greater than that of *Microcystis* despite the fact that these were *Microcystis* blooms. These contaminants could therefore have had a skewing effect on the abundance results.

Detecting the occurrence blooms is important for mitigation, but the ability to distinguish toxic from non-toxic blooms is also critically important. The *mcy* genes code for the microcystin toxin that makes *Microcystis* blooms so harmful. We estimated the abundance of reads aligning to the *Microcystis mcy*-family genes as part of this objective. We used the reads from our initial set of pre-bloom samples for this analysis. We aligned all the reads classified to the *Microcystis* genus against a reference consisting of all annotated *Microcystis* genomes using the aligner minimap2 (Li 2018), and then filtering the alignments to those intersecting the genomic intervals annotated as containing *mcy*-family genes using bedtools (Quinlan and Hall 2010). We saw four hits in the full run pre-bloom sample from GLERL WE13, two to *mcyD*, and one each to *mcyC*, and *mcyE*. The WE02 sample produced no hits at all.

We also subsampled the pre-bloom sequencing runs using the timestamps added by the MinION software for all reads as they are sequenced. We generated a table of read IDs and their time stamps, which were then joined back to the WIMP classification results. We binned the reads into cumulative one-hour and four-hour buckets based on the time of sequencing.

Based on the accumulation data, we can conclude that for this protocol, the majority of the reads for all samples were sequenced within 24 hours of beginning the run. Having subsampled the reads, we then evaluated the length of run time necessary to accurately approximate the range of relative abundances of the genus *Microcystis*, and the species and strains of that that we observed in our samples. By 12 hours, all but the rarest taxa are detectable, and the overall relative abundance of taxa appears consistent from that point forward. By 24 hours, the rarer taxa are detected and the pattern of relative abundances changes very little. Similarly, the relative abundance of both the genus *Microcystis* and the species *Microcystis aeruginosa* converge very close to their final values by 16 hours into the run, with almost no deviation after 24 hours. Our preliminary conclusion is that a 16 to 24 hour run (equivalent to multiplexing two to three samples on a flow cell) is sufficient to capture *Microcystis* abundance reliably for water samples of this type. However, there are significant caveats to this conclusion based on two factors. First, the contaminants that we discovered may have affected our abundance estimates, especially relative abundance. Second, the low yields of DNA extracted from our samples mean that our overall read counts were lower than expected for this MinION protocol.

We next sequenced samples drawn from bloom conditions from the same GLERL stations during the same year as our pre-bloom samples. Downstream processing and taxonomic classification were performed using the same MinKNOW and WIMP pipelines as used for the initial pre-bloom samples. We multiplexed the bloom and pre-bloom samples on our two remaining flow-cells using the ONT Rapid Barcoding Kit 96 (SQK-RBK110.96). Two bloom samples and two pre-bloom samples (one of each from each GLERL) were sequenced on each flow cell to avoid the possibility of batch effects. Using the per-taxon read counts generated for each sample with the WIMP pipeline, we then performed a differential abundance analysis using DESeq2 (Love, Huber, and Anders 2014). This analysis passed an essential quality assurance check, in that all taxa of *Microcystis* were significantly more abundant in the bloom condition than in the pre-bloom condition. However, the range of different taxa that are differentially abundant suggests the possibility of a pre-bloom taxon profile that might be used as features in a predictive model for forecasting blooms. Taxa such as *Methylobacterium* and *Stenotrophomonas* that are significantly more abundant pre-bloom compared to during the bloom might be of particular interest as indicator taxa, for which a shift in abundance might presage a HAB. As with the pre-bloom samples, we also investigated the abundance of the *mcy*-family genes for the bloom samples generated as part of this objective. Reads classified by WIMP as belonging *Microcystis aeruginosa* were aligned against a reference set of all annotated *Mycrocystis* genome sequences, and then the alignments were then intersected with the annotated *mcy* gene positions. While pre-bloom samples in few hits (four reads mapping to *mcyC*, *mcyD*, and *mcyE* in one full-run sample) we did observed more hits in the bloom-drawn samples. The bloom samples in GLERL WE02 contained hits to all 10 *mcy* genes. By contrast, in station WE13, which suffered a much less significant bloom in 2019 (the year of sampling) had reads mapping only to some members of the gene family.

## 5 Acknowledgements, Author Contributions & Competing Interests

The authors would like to express sincere gratitude and acknowledge Jessica Bouchet, David Russell, and Hunter Baylous for assisting with laboratory work on this project, and Dr. Stephanie Guertin for consultation on design and execution.

AK performed sequencing and conducted all data analysis. All authors contributed to study design, results interpretation, and drafting and reviewing the manuscript.

This material is based upon work supported by the SBIR Program within the NOAA Technology Partnerships Office under Grant No. NA21OAR0210481.

AK and WG are employees of Elder Research, recipient of the NOAA SBIR grant that funded the collection of this data. ST is an employee of Signature Science, LLC.

